# The effects of oral supplementation of Japanese sake yeast on anxiety, depressive-like symptoms, oxidative stress, and BDNF changes in chronically stressed adolescent rats

**DOI:** 10.1101/2022.09.14.507940

**Authors:** Motahareh Haghipanah, Maryam Saadat, Asal Safarbalou, Thomas Budde, Wael Mohamed, Elham Sadat Afraz, Nasrollah Moradikor

## Abstract

Chronic stress during the pre-pubertal period has adverse effects in developing neuropsychiatric disorders such as depression and anxiety. The administration of supplements with antioxidant properties may alleviate depression and anxiety behavior. This study investigated the effects of oral supplementation of Japanese sake yeast on anxiety, depressive-like symptoms, oxidative stress, and changes in brain-derived neurotropic factor (BDNF) in adolescence rats subjected to chronic stress.

In order to assess the effects of chronic stress, adolescent rats were grouped into one non-stressed control group (non-stress) and four different experimental groups. The other animals were subjected to stress and orally received normal saline (Control-stress), 15 mg/kg yeast (Stress-15), 30 mg/kg yeast (Stress-30) and 45 mg/kg yeast (Stress-45). Anxiety and depression-like behavior, BDNF levels, and oxidative stress markers were evaluated.

The rats exposed to stress exhibited anxiogenic and depression-like behavior as well as lower levels of BDNF and higher levels of oxidative markers compared with non-stressed rats (*P*<0.05). However, the oral supplementation of sake yeast decreased anxiogenic and depression-like behavior and oxidative indices, and also increased BDNF levels compared to stressed rats treated with saline in a dose-dependent manner (*P*<0.05).

In sum, stress caused anxiety and depression behavior, increased oxidative indices, and reduced BDNF levels while sake yeast alleviated adverse effects of stress on anxiety and depression behaviors, decreased oxidative markers, and increased BDNF levels.

## Introduction

Chronic stress in the pre-pubertal age is known to have adverse effects in progressing neuropsychiatric disorders in adulthood [1]. Adolescence is known as a sensitive age in which the development of psychopathology may occur [2]. Indeed, most anxious and depressive episodes occur during this period [3]. Stressors and immune challenges increase vulnerability to neurodegeneration during the adolescence period [4]. Stress raises anxiety and depression in adolescences because it interferes with neuronal function and morphology in corticolimbic structures such as amygdala and other emotional regulatory parts in the brain [5]. The association between stress and development of psychological disorders such as depression has been shown [6–8]. Stress has been reported to be responsible for decreased hippocampal neurogenesis, which is associated with depression. [9]. Indeed, chronic stress lowers the levels of brain-derived neurotropic factor (BDNF) that increases vulnerability to depression [10]. On the other hand, the brain is a vulnerable organ to oxidative damage owing to its high polyunsaturated fatty acids and low antioxidant capacity [11, 12]. Increased malondialdehyde (MDA) is considered to be the key factor for oxidative stress in mood disorders [13]. In addition, the involvement of oxidative stress and antioxidant enzymes like superoxide dismutase (SOD) and glutathione peroxidase (GPx) in anxiety-like behavior in rodents has been reported [14]. Various agents are utilized for the treatment of depression and anxiety, but they may have diverse side effects. It is, therefore, necessary to find safe agents with multifunctional efficiency.

Sake yeast, which is used to make Japanese rice wine, is utilized in the production of alcoholic beverages since ancient times [15]. It is known to have antioxidant activity in an animal model with chronic diabetic conditions. [16]. Indeed, sake is the national alcohol beverage of Japan and is extensively consumed because of its refreshing taste and elegant flavor [17]. Our literature search did not reveal any study investigating the effects of sake yeast on depression and other mood disorders yet. We hypothesized that yeast sake may improve signs of depression and anxiety owing to its antioxidant properties. This preliminary study aimed to investigate the effects of oral supplementation of Japanese sake yeast on anxiety, depressive-like symptoms, oxidative stress, and BDNF changes in adolescence rats subjected to chronic stress.

## Materials and methods

### Sake yeast

Sake yeast powder (GSP6) was provided from Lion Corporation (Odawara-shi, Japan).

### Animals

All care procedures for animals were in agreement with the Ethical Committee of International Center for Intelligent Research (ICIR-2021-186672). In the current study, 60 rats in five groups were studied. Four groups were stressed, and one group was not submitted to stress and was considered a non-stress group. Three of the stressed groups received sake yeast powder (15 mg/kg yeast, Stress-15; 30 mg/kg yeast, Stress-30; 45 mg/kg yeast, Stress-45) and one normal saline (Control-stress). Sake yeast doses were selected based on previous studies [16]. Animals had free access to water and food; their experimental environment was continually monitored for temperature and water.

### The induction of stress

Stress was induced as reported previously [18, 19]. In summary, animals were submitted to restraint stress (2 h/day for 10 consecutive days) in a clear polyethylene cylinder. To immobilize, the cylinder size was regulated based on the size of the pup rats and had a hole in the front of the cylinder for improved breathing.

### Anxiety and depression-like behavior tests

#### Open-field test (OFT)

The OFT was conducted to assess anxiety as reported by others [1]. Briefly, behavioral responses in time were investigated in a dark area (72×72×45 cm) for 20 minutes.

#### Elevated plus maze (EPM)

The EPM was utilized to assess anxiety as reported by others [1]. In summary, behavioral responses were assessed in apparatuses consisted of two open arms (50 cm × 10 cm) and two enclosed arms (50 cm × 10 cm, surrounded by 40-cm high wooden walls), raised 50 cm above the floor.

#### Force swimming test (FST)

The FST was performed based on previous studies [20]. In short, the animals were kept in a cylindrical swimming tank (50 cm high, 25 cm diameter) that was filled with 25 °C water for 15 minutes and was dried and cleaned thereafter. Rats were placed into the swimming tank for 5 minutes after 24 hours. In the current study, we investigated immobility, swimming, and climbing. The increased immobility and decreased swimming and climbing were considered depressive behavior.

### The assessment of BDNF and antioxidant-associated factors

The animals were decapitated, and the whole prefrontal cortex (PFC) was dissected and then immediately frozen at −80°C. Tissue was homogenized in cold lysis buffer, and BDNF was assessed by ELISA kits (Hangzhou Eastbiopharm Co., LTP) following manufacturer instruction. Brain sections were frozen, homogenized, and investigated for MDA by MDA ELISA kits (Hangzhou Eastbiopharm Co., LTP) following manufacturer instruction. SOD and GPx activities in brain were also assessed by ZellBio kits (Zellbio GmbH, Ulm, Germany) following manufacturer instruction.

### Data analysis

Data were checked for normality using the Shapiro-Wilk-Test. Having found the normal distribution, we used parametric tests for determining statistical significant differences. Data were analyzed using Graph Pad Prism Software (version of 6.07), and mean ± SD was reported.

## Results

Figure 1 depicts the results for oral supplementation of sake yeast on anxiety-like behavior in stressed rats. The induction of stress significantly decreased the number of visits, center time, and total distance (control stress and non-stress rats). Compared to the control stress group, supplementation of sake yeast significantly increased the number of visits, center time, and total distance in a dose-dependent manner. In sum, stress induced anxiety was alleviated by the effects of supplementation of sake yeast.

**Figure 1.**
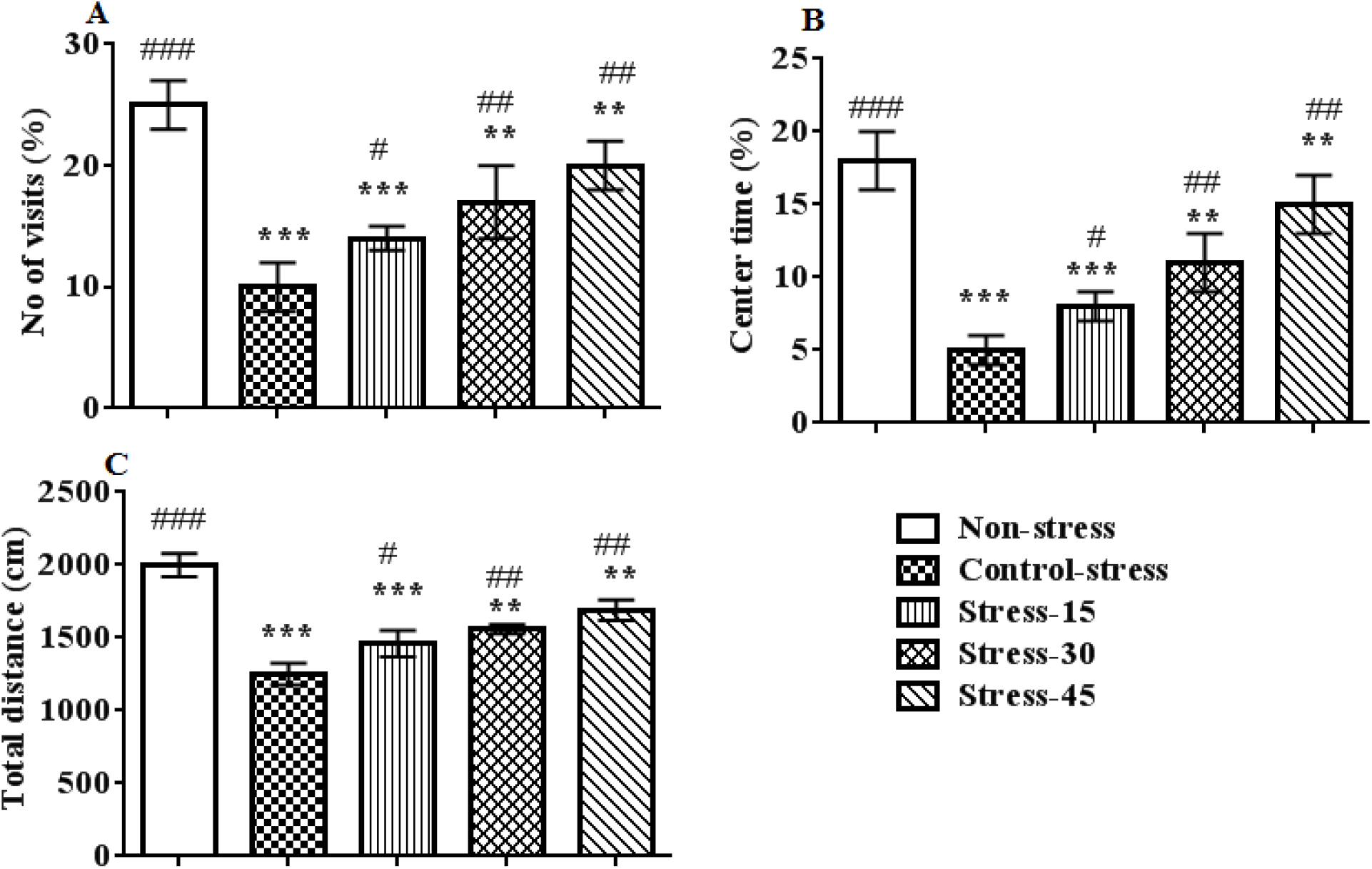
The results for oral supplementation of sake yeast on anxiety-like behaviors in stressed rats. Superscript symbols * and # indicate significant differences between non-stress and the other groups and between control-stress and the other groups, respectively.

Figure 2 shows the results of oral supplementation of sake yeast on elevated plus maze behavior in stressed rats. The control-stress group spent less time in the open arm compared with non-stressed group (P=0.0001). The administration of sake yeast significantly increased the time and number of open arm visits compared with those of the control-stressed group. In sum, stress caused anxiety but supplementation of sake yeast alleviated it.

**Figure 2.**
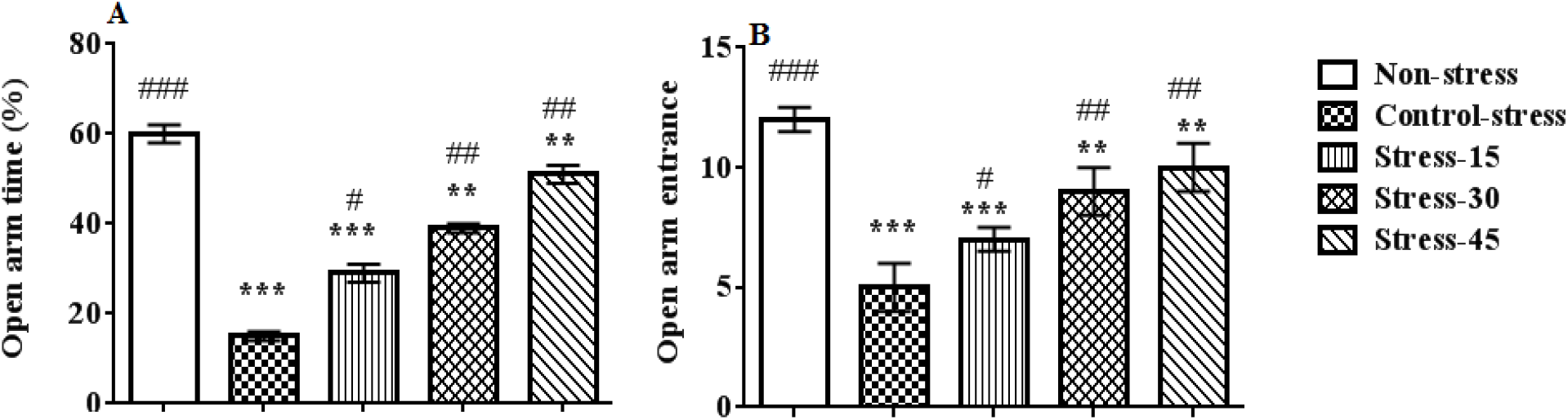
The results for oral supplementation of sake yeast on elevated plus maze behaviors in stressed rats. Superscript symbols * and # indicate significant differences between non-stress and the other groups and between control-stress and the other groups, respectively.

Figure 3 shows the results of oral supplementation of sake yeast on depression-like behavior in stressed rats. Our findings showed that stress decreased the duration of swimming while phases of immobility were prolonged, highlighting the induction of depression. The supplementation of sake yeast increased swimming duration and decreased immobility duration in a dose-dependent manner (P=0.0001). In short, stress induced depression was alleviated by the supplementation of sake yeast.

**Figure 3.**
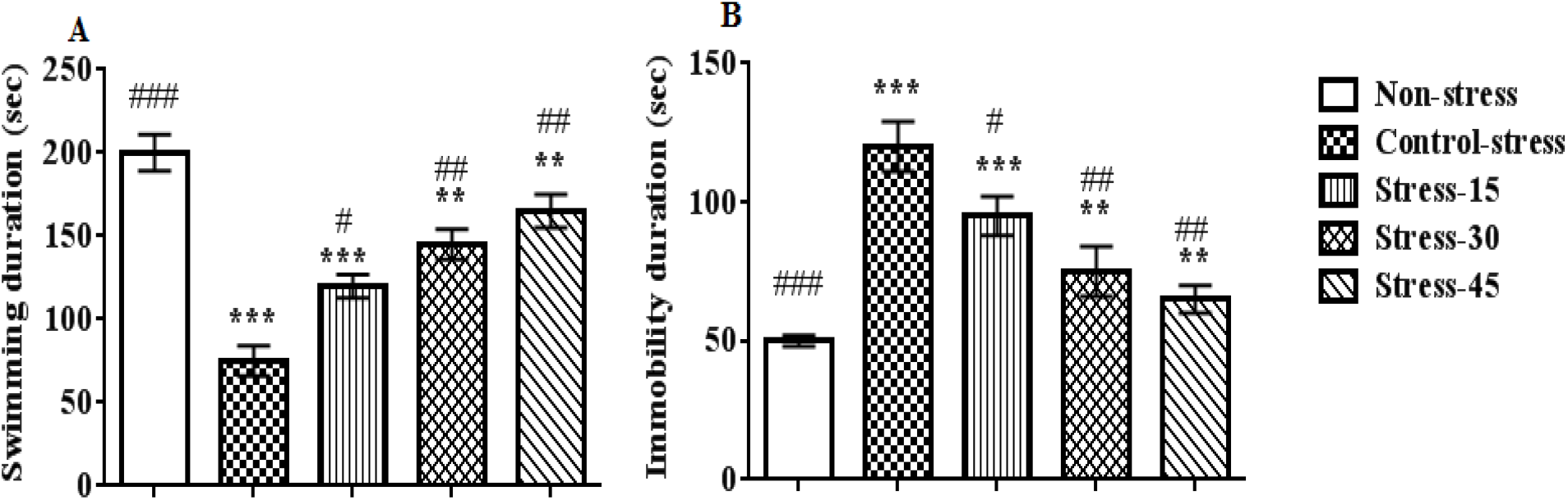
The results for oral supplementation of sake yeast on depression-like behaviors in stressed rats. Superscript symbols * and # indicate significant differences between non-stress and the other groups and between control-stress and the other groups, respectively.

Figure 5 depicts our findings for oral supplementation of sake yeast on prefrontal cortex BDNF levels in stressed rats. Non-stressed rats had high BDNF levels compared to control-stress. Furthermore, the oral supplementation of sake yeast dose-dependently increased prefrontal cortex BDNF levels compared with control-stress (P=0.0001).

**Figure 4.**
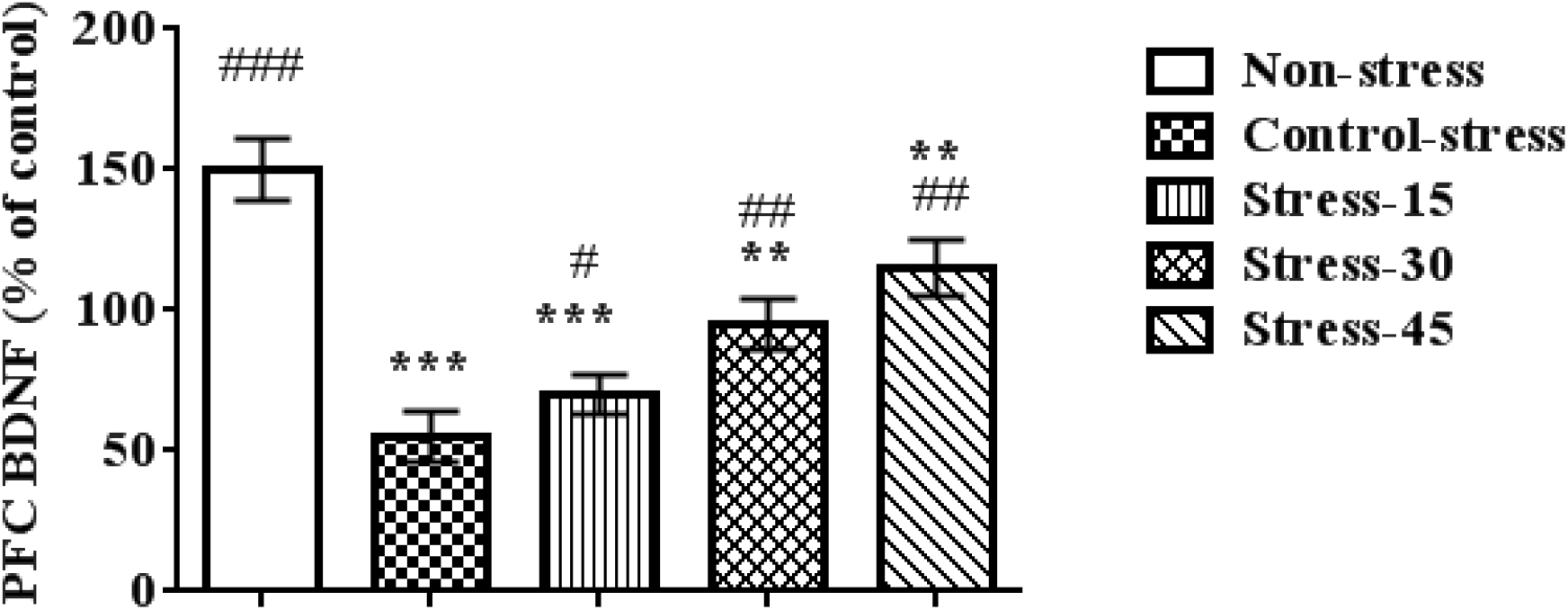
The results for oral supplementation of sake yeast on prefrontal cortex BDNF levels in stressed rats. Superscript symbols * and # indicate significant differences between non-stress and the other groups and between control-stress and the other groups, respectively.

**Figure 5.**
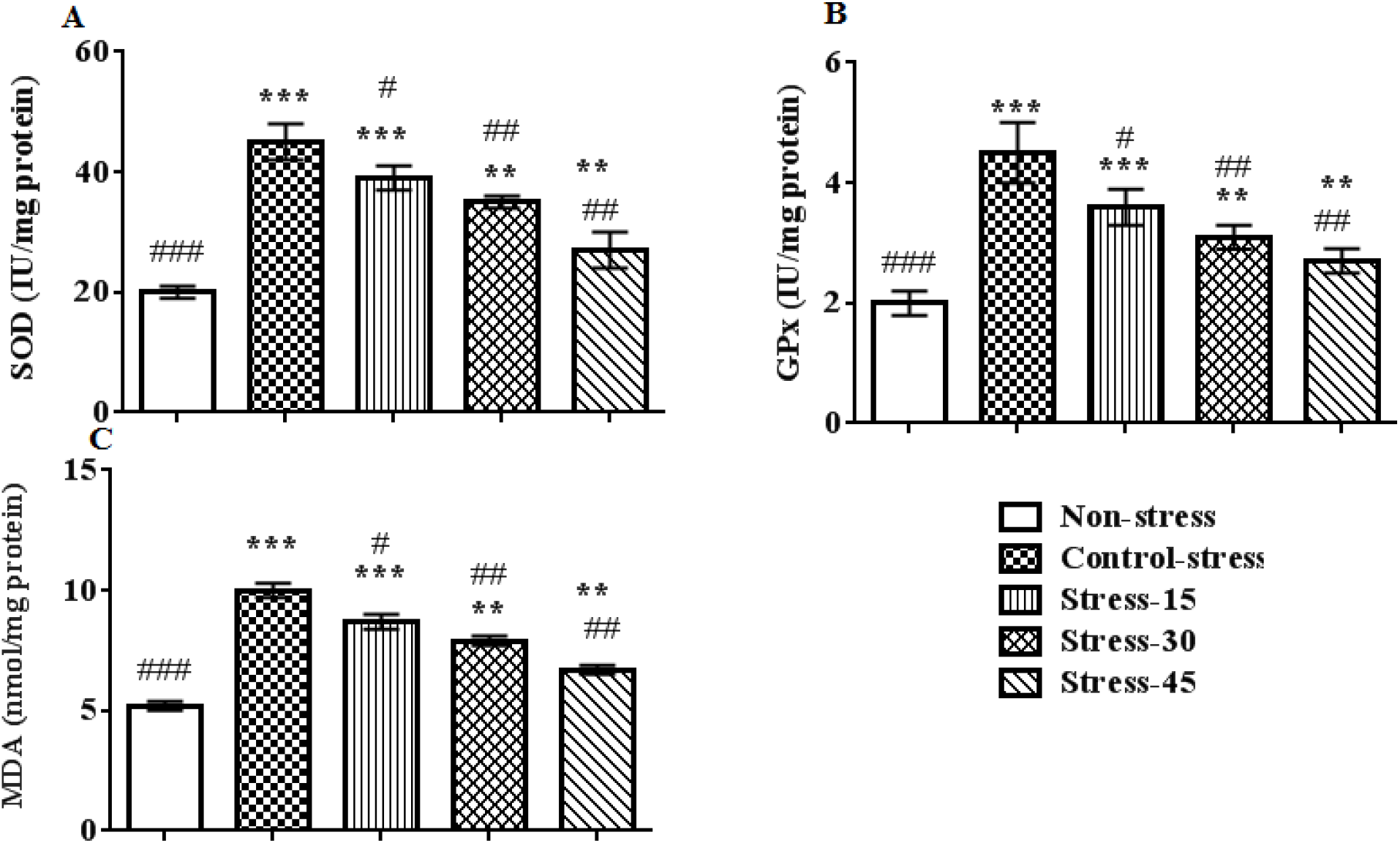
The results of oral supplementation of sake yeast on activities of antioxidant enzymes and MDA levels in stressed rats. Superscript symbols * and # indicate significant differences between non-stress and the other groups and between control-stress and the other groups, respectively.

Figure 5 shows the results for oral supplementation of sake yeast on activities of antioxidant enzymes and MDA levels in stressed rats. Our findings showed that induction of stress significantly increased the activities of antioxidant enzymes and MDA levels (P=0.0001). The administration of sake yeast dose-dependently decreased activities of antioxidant enzymes and MDA levels (P<0.001).

## Discussion

Our findings showed that stress during adolescence was parallel with anxiogenic and depression-like behavior in adulthood. Based on our findings, the animals exposed to stress spent less time in open arms and had fewer open arm entries compared with non-stressed rats. In addition, center time and center distance were lower in stressed rats compared with non-stressed rats. Moreover, we observed anxiety and depression behavior after the induction of stress in non-treated rats. The results for the effects of stress on depression are in line with previous studies that showed stress could induce depression and anxiety [8, 21, 22]. Stress increases depression and anxiety via increased levels of corticosterone [1, 18]. Importantly, our findings showed that sake yeast alleviated the effects of stress. The mechanism of sake yeast in decreasing depression and anxiety could be attributed to its effects on BDNF and antioxidant factors, as will be discussed below.

Our findings showed that stress and sake yeast had opposing effects on brain BDNF levels. While stress decreased the BDNF level, sake yeast increased it. The BDNF-TrkB pathway is beneficial in regulating mood and emotional behaviors [23]. It was reported that intra-hippocampal infusions of BDNF improves behavioral responses in an animal model of depression [24]. It has been reported that exposure to stress decreases BDNF expression [25, 26], which is consistent with our findings. Sake yeast increased BDNF levels, thus corroborating its effects in decreasing anxiety and depression. In literature, studies investigating the effects of sake yeast on brain BDNF levels were not found.

The results further showed that stress increased activity of antioxidant enzymes and MDA levels. Brain cells are susceptible to oxidative damage because the brain has significant oxygen circulation, and polyunsaturated fatty acids expose it to lipid peroxidation. [27]. The brain typically responds to stressors via increased antioxidant enzymes [28]. Increased lipid peroxidation is closely related to increased MDA in the brain [29]. MDA is considered to be an appropriate biomarker for oxidative stress in mood disorders [13]. Sake yeast modulated antioxidant enzyme activity and MDA levels, which is consistent with our previous study [16]. It was reported that sake yeast modulated antioxidant enzyme activities via lowering hyperglycemia, inflammation, and oxidative stress. In sum, we found that sake yeast acts as an antioxidant and decreases antioxidant activities in the brain. It also protects brain from damages and stressors that cause depression and anxiety.

## Conclusions

In conclusion, stress increases anxiety and depression-like behavior, activity of antioxidant enzymes, and it decreases BDNF levels in adolescent rats. The oral supplementation of sake yeast significantly decreases anxiety and depression-like behavior, activity of antioxidant enzymes, and it increases BDNF levels. Oral supplementation of sake yeast alleviated depression and anxiety behavior via improving the antioxidant status and BDNF levels. Currently, the relevance of our findings from animal studies to the human situation is not clear. However, the beneficial effects of sake yeast supplementation on the quality of sleep in humans indicate that our preliminary study is promising and opens a new window for future studies.

## Authors’ contributions

All authors contributed toward data analysis, drafting and revising the paper and agreed to be responsible for all the aspects of this work.

